# Inhibition of cleavage of human complement component C5 and the R885H C5 variant by two distinct high affinity anti-C5 nanobodies

**DOI:** 10.1101/2023.02.22.529391

**Authors:** Eva M. Struijf, Karla I De la O Becerra, Maartje Ruyken, Fleur van Oosterom, Danique Y. Siere, Dani A. C. Heesterbeek, Edward Dolk, Raimond Heukers, Bart W. Bardoel, Piet Gros, Suzan H.M. Rooijakkers

**Affiliations:** Medical Microbiology, University Medical Center Utrecht, Utrecht University, Utrecht, the Netherlands; Structural Biochemistry, Bijvoet Centre for Biomolecular Research, Department of Chemistry, Faculty of Science, Utrecht University, Utrecht, the Netherlands; QVQ Holding BV, Utrecht, The Netherlands

**Keywords:** complement system, innate immunity, nanobody, single-domain antibody (sdAb nanobody), inhibitor, protein structure, structural model, inflammatory diseases

## Abstract

The human complement system plays a crucial role in immune defense. However, its erroneous activation contributes to many serious inflammatory diseases. Since most unwanted complement effector functions result from C5 cleavage, development of C5 inhibitors, such as clinically approved monoclonal antibody Eculizumab, are of great interest. In this study, we developed and characterized two anti-C5 nanobodies, UNbC5-1 and UNbC5-2. Using surface plasmon resonance (SPR), we determined a binding affinity of 120 pM for UNbC5-1 and 8 pM for UNbC5-2. Competition experiments determined that the two nanobodies recognize distinct epitopes on C5. Both nanobodies efficiently interfered with C5 cleavage in a human serum environment, as they prevented red blood cell lysis via membrane attack complexes (C5b-9) and the formation of chemoattractant C5a. The cryo-EM structure of UNbC5-1 and UNbC5-2 in complex with C5 revealed that the binding interfaces of UNbC5-1 and UNbC5-2 overlap with known complement inhibitors Eculizumab and RaCI3, respectively. UNbC5-1 binds to the MG7 domain of C5, facilitated by a hydrophobic core and polar interactions, and UNbC5-2 interacts with the C5d domain mostly by salt bridges and hydrogen bonds. Interestingly, UNbC5-1 potently binds and inhibits C5 R885H, a genetic variant of C5, that is not recognized by Eculizumab. Altogether, we identified and characterized two different, high affinity nanobodies against human C5. Both nanobodies could serve as diagnostic and/or research tools to detect C5 or inhibit C5 cleavage. Furthermore, the residues targeted by UNbC5-1 hold important information for therapeutic inhibition of different polymorphic variants of C5.

## Introduction

The complement system is an important part of the human innate immune system that is involved in many different biological processes. During embryonal tissue development, it plays a crucial role in cell differentiation and proliferation (1–3). Later in life, complement is best known for its role in clearing pathogens during infection, but also remains involved in maintaining homeostasis, by removing apoptotic cells and immune complexes (4). Although the complement system is tightly regulated, unwanted complement activation is the root cause of a variety of diseases, including paroxysmal nocturnal hemoglobinuria (PNH), atypical hemolytic uremic syndrome (aHUS), and age-related macular degeneration (AMD). Furthermore, erroneous complement activity contributes in more prevalent complex multifactorial diseases like systemic lupus erythematosus (SLE), Alzheimer’s disease, and COVID-19 (5).

The complement system can be activated via three distinct pathways: the classical pathway (CP), the lectin pathway (LP), and the alternative pathway (AP). Together, these pathways result in three major effector functions: [1] opsonization, [2] chemo-attraction of immune cells, and [3] direct target cell lysis (6). A central complement component is C5, a 190 kDa protein, consisting of two disulfide linked chains (alpha 115 kDa and beta 75 kDa), which circulates in blood at a concentration of ±75 μg/ml (7). During complement activation, C5 convertases (C4b2bC3b and C3bBbC3b) are formed on the target surface, which cleave native C5 molecules in C5a (8 kDa) and C5b (181 kDa). While C5a is released in the supernatant where it functions as a chemoattractant, C5b interacts with complement proteins C6, C7, C8, and multiple copies of C9, forming the membrane attack complex (MAC) (6).

Since multiple (unwanted) complement effector functions result from the cleavage of C5, many complement inhibitors have been identified and developed to target this particular molecule. Human pathogens have evolved immune evasion proteins to specifically block the cleavage of C5. For example, *Staphylococcus aureus* expresses SSL7, a protein that potently binds and inhibits cleavage of human C5 (8). Furthermore, ticks (*Ornithodoros moubata, Rhipicephalus pulchellu*s, and *Rhipicephalus appendiculatus*) contain C5 inhibiting proteins OmCI (9), CirpT (10), and RaCI3 (11), respectively, in their saliva.

In addition to pathogen-derived C5 inhibitors, a wide variety of C5-targeting molecules, including monoclonal antibodies have been developed (12, 13). Of these, the best known is the monoclonal antibody Eculizumab, that is FDA approved, and which binds to C5 and prevents its cleavage by blocking the interaction of C5 with the C5 convertases (14). After the introduction of Eculizumab into the clinic, a genetic polymorphism in C5 (C5 R885H) was identified in a group of poor responders (15). The structural model of Eculizumab and C5 revealed that R885 in C5 and P101 in Eculizumab form a crucial interaction for binding and inhibiting C5 (16, 17).

In this study, we identify and characterize two llama-derived nanobodies that specifically bind and inhibit human complement C5. They bind to C5 with picomolar affinities on two distinct epitopes. Interestingly, one of these nanobodies has an overlapping epitope with Eculizumab and is still able to inhibit C5 R885H.

## Results

### Identification of two C5-targeting nanobodies that interfere with complement

To develop nanobodies targeting C5, llamas were immunized with recombinant human C5 and nanobody phage libraries were generated. After two rounds of phage display panning, ∼200 clones were produced in the periplasm of *Escherichia coli* and unpurified periplasmic fractions were used to screen for clones that bind C5 in ELISA. Based on these ELISA signals and nanobody sequences, two positive clones were selected, denoted UNbC5-1 and UNbC5-2. These two nanobodies were significantly distinct, with having only 30% amino acid sequence similarity in their complementarity determining regions (CDR) (**Fig. 1A**). Next, UNbC5-1 and UNbC5-2 were purified to assess their ability to block complement activity in human serum. Using a CP hemolysis assay (18), we identified UNbC5-1 and UNbC5-2 as complement inhibitors (**Fig. 1B**). Both nanobodies blocked complement-mediated lysis in a dose-dependent manner, with IC50 values that are in the same range as C5 inhibitor RaCI3 (**Table I**). Eculizumab was roughly 3-15 times more potent in blocking C5 cleavage than UNbC5-1 and UNbC5-2. This might partially be explained by the fact that Eculizumab has two antigen-binding domains per molecule and the nanobodies have only one.

**FIGURE 1:**
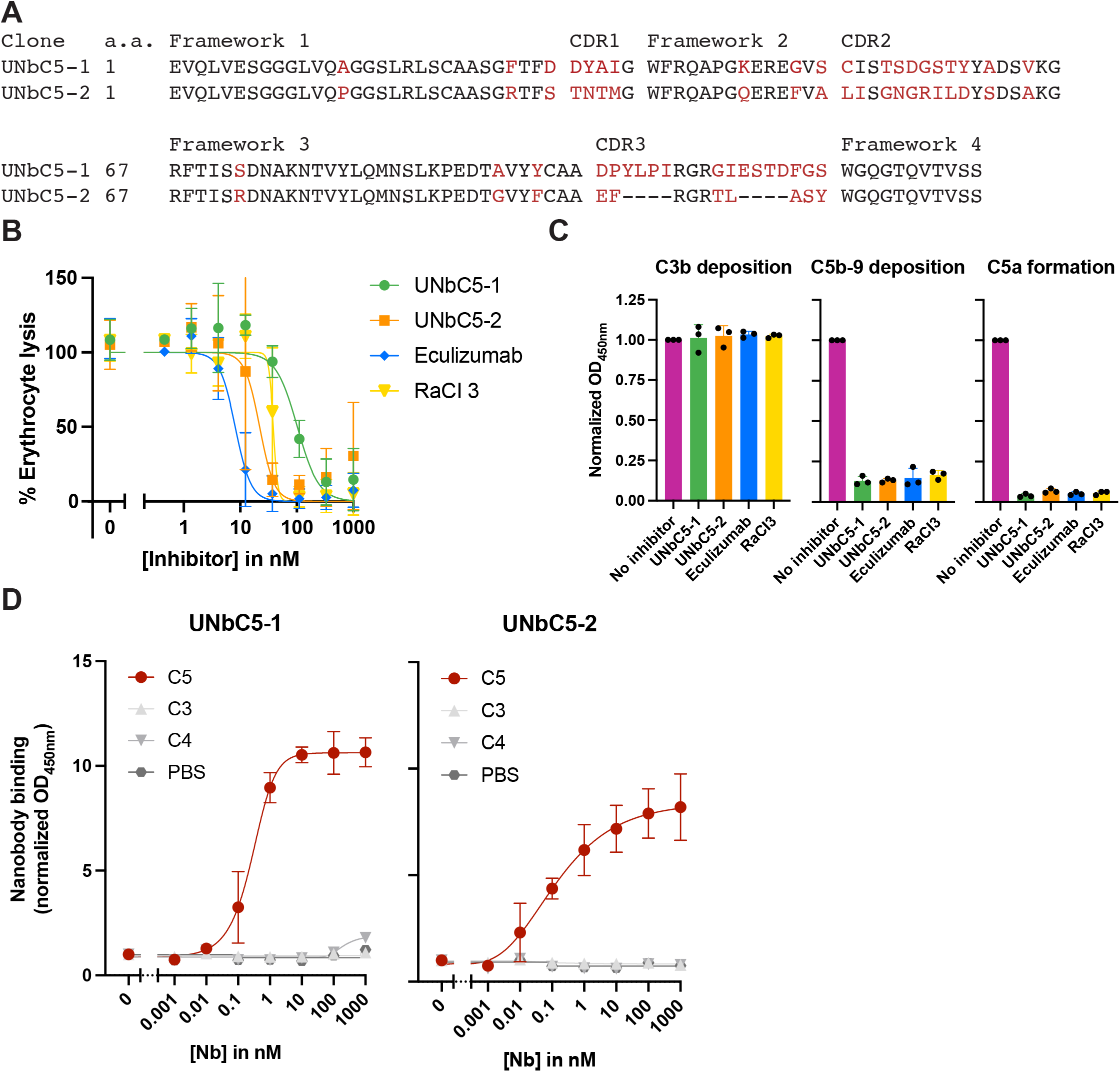
Identification of two C5-targeting nanobodies that interfere with complement. (A) Protein sequence alignment of UNbC5-1 and UNbC5-2. Frameworks 1-4 and CDRs 1-3 are indicated. Conserved amino acids are indicated in black and unique amino acids are indicated in red. (B) CP mediated hemolysis of antibody-coated sheep erythrocytes incubated with 2.5% normal human serum and a titration of our nanobodies UNbC5-1 and UNbC5-2 and known complement inhibitors RaCI3 and Eculizumab. The OD405 values of the supernatants were measured, the % erythrocyte lysis was calculated using the 0% lysis (buffer) and 100% lysis (MilliQ) control samples. (C) CP complement activation on an IgM-coated microtiter plate, incubated with 4% human serum in the presence of buffer (no inhibitor) or 1000 nM inhibitor. Deposition of complement activation products C3b and C5b-9 on the plate was measured using specific anti-C3 and anti-C5b-9 antibodies and HRP-coupled secondary antibodies, at OD450. C5a formation was measured by adding the supernatant of the reaction to a microtiter plate coated with C5a capture antibodies and C5a was detected with a specific C5a detection antibody and HRP-coupled secondary antibodies, at OD450. Data points were normalized to the maximum levels of C3b and C5b-9 deposition and to the maximum levels of C5a formation when no inhibitor was added. (D) Nanobody binding to complement protein C5 and not homologs C3 and C4. Microtiter plates were coated with complement proteins and incubated with increasing concentrations of UNbC5-1 and UNbC5-2. Nanobody binding was measured using polyclonal rabbit-anti-VHH QE19 antibodies and donkey-anti-rabbit-HRP antibodies, at an OD of 450 nm. Coating with PBS was taken along as a negative control. Data information: (A) sequences were aligned using T-coffee (44). (B-D) Data represent mean ± SD of 3 individual experiments (B & D) curves were fitted and IC50 and EC50 values were obtain.

**Table I.** IC50 and EC50 values for UNbC5-1, UNbC5-2, Eculizumab and RaCI3 for inhibiting complement activity and binding complement C5.

|  | IC<sub>50</sub> (nM) | EC<sub>50</sub> (nM) | EC<sub>50</sub> (nM) |
| --- | --- | --- | --- |
|  | CP | C5 WT | C5 R885H |
| UNbC5-1 | 137 ± 83 | 0.38 ± 0.2 | 1.5 ± 0.4 |
| UNbC5-2 | 24 ± 15 | 0.12 ± 0.01 | 22.43 ± 17.0 |
| Eculizumab | 9 ± 4 | 0.049 ± 0.013 | ND |
| RaCI3 | 42 ± 8 | - | - |

Binding and inhibition curves were fitted using the GraphPad Prism 9.3.0 function “[inhibitor] vs. normalized response – variable slope” & “Asymmetrical Sigmoidal, 5PL, X is concentration”. IC<sub>50</sub> and EC<sub>50</sub> values were calculated for 3 individual experiments, using the fitted curves and SD were determined.

To confirm that UNbC5-1 and UNbC5-2 block complement at the level of C5, we assessed their capacity to block different steps of the complement reaction. Using an ELISA-based complement activity assay (19), we observed that neither UNbC5-1 nor UNbC5-2 affect the initial steps of complement activation since there is no inhibition of C3b deposition on IgM-coated microtiter plates that were incubated with serum (**Fig. 1C**). However, UNbC5-1 and UNbC5-2 can prevent the formation of deposited C5b-9 complexes and the release of C5a into the supernatant. Thus, we show that UNbC5-1 and UNbC5-2 block complement activity at the level of C5 cleavage. Consistent with this finding, UNbC5-1 and UNbC5-2 could also prevent complement-mediated hemolysis by the alternative pathway (**SFig. 1A**).

To assess whether UNbC5-1 and UNbC5-2 are indeed specific for C5, we studied the binding to complement proteins C3 and C4, which share sequence and structural homology with C5 (20). In ELISA, UNbC5-1 and UNbC5-2 showed a dose-dependent binding to C5, while no cross reactivity with C3 or C4 was observed (**Fig. 1D**). UNbC5-1 and UNbC5-2 bind C5 with EC50 values of 383 ± 220 and 122 ± 14 pM, respectively (**Table I**). Altogether these data show that UNbC5-1 and UNbC5-2 are C5-specific nanobodies that can interfere with complement activity.

### UNbC5-1 and UNbC5-2 bind C5 with picomolar affinity and recognize distinct epitopes

Next, we used surface plasmon resonance (SPR) to determine the affinities of UNbC5-1 and UNbC5-2 for C5. Briefly, biotinylated nanobodies were coupled to streptavidin coated biosensors and a kinetic titration with C5 was performed to determine k_on_ rates (**Table II**). Next, a dissociation step was performed to measure k_off_ rates and K_D_ values were calculated. UNbC5-1 binds C5 with a K_D_ of 120 pM (**Fig. 2A**) and UNbC5-2 binds C5 with a kD of 8 pM (**Fig. 2B**). As a reference, we took along Eculizumab and measured a C5 binding affinity of 104 pM, which is in the same range as its published affinity (17.6-120 pM) (**SFig. 3)** (16, 21, 22). To assess whether the two nanobodies bind overlapping epitopes, we performed competition experiments by ELISA. C5-coated microtiter plates were incubated with Myc-labeled UNbC5-1 (UNbC5-1-Myc) in the presence or absence of unlabeled UNbC5-2. After washing, binding of UNbC5-1-Myc to C5 was quantified. We observed that the presence of UNbC5-2 did not affect binding of UNbC5-1-Myc to C5 (**Fig. 2C**). As a control, we showed that UNbC5-1 was able to compete with UNbC5-1-Myc. These data indicate that UNbC5-1 and UNbC5-2 can bind C5 simultaneously and not compete for the same epitope.

**Table II.** Binding affinities of UNbC5-1, UNbC5-2 and Eculizumab for complement C5.

| Nanobody | $k_{\text{on}}$ ( $\text{M}^{-1}\text{s}^{-1}$)$\times 10^5$ | $k_{\text{off}}$ ($\text{s}^{-1}$)$\times 10^{-6}$ | $K_{\text{D}}$ (pM) |
| --- | --- | --- | --- |
| UNbC5-1 | $2.713 \pm 0.002$ | $3.254 \pm 0.06$ | $119.90 \pm 0.20$ |
| UNbC5-2 | $2.937 \pm 0.008$ | $0.227 \pm 0.03$ | $7.74 \pm 0.09$ |
| Eculizumab | $3.661 \pm 0.003$ | $3.810 \pm 0.10$ | $104.00 \pm 0.40$ |

Nanobody and Eculizumab affinities were determined by SPR using C5 as analyte and nanobodies and Eculizumab as a ligand. All measurements were performed in duplicate.
Kinetics were determined using Scrubber 2.0 (BioLogic Software).

**FIGURE 2:**
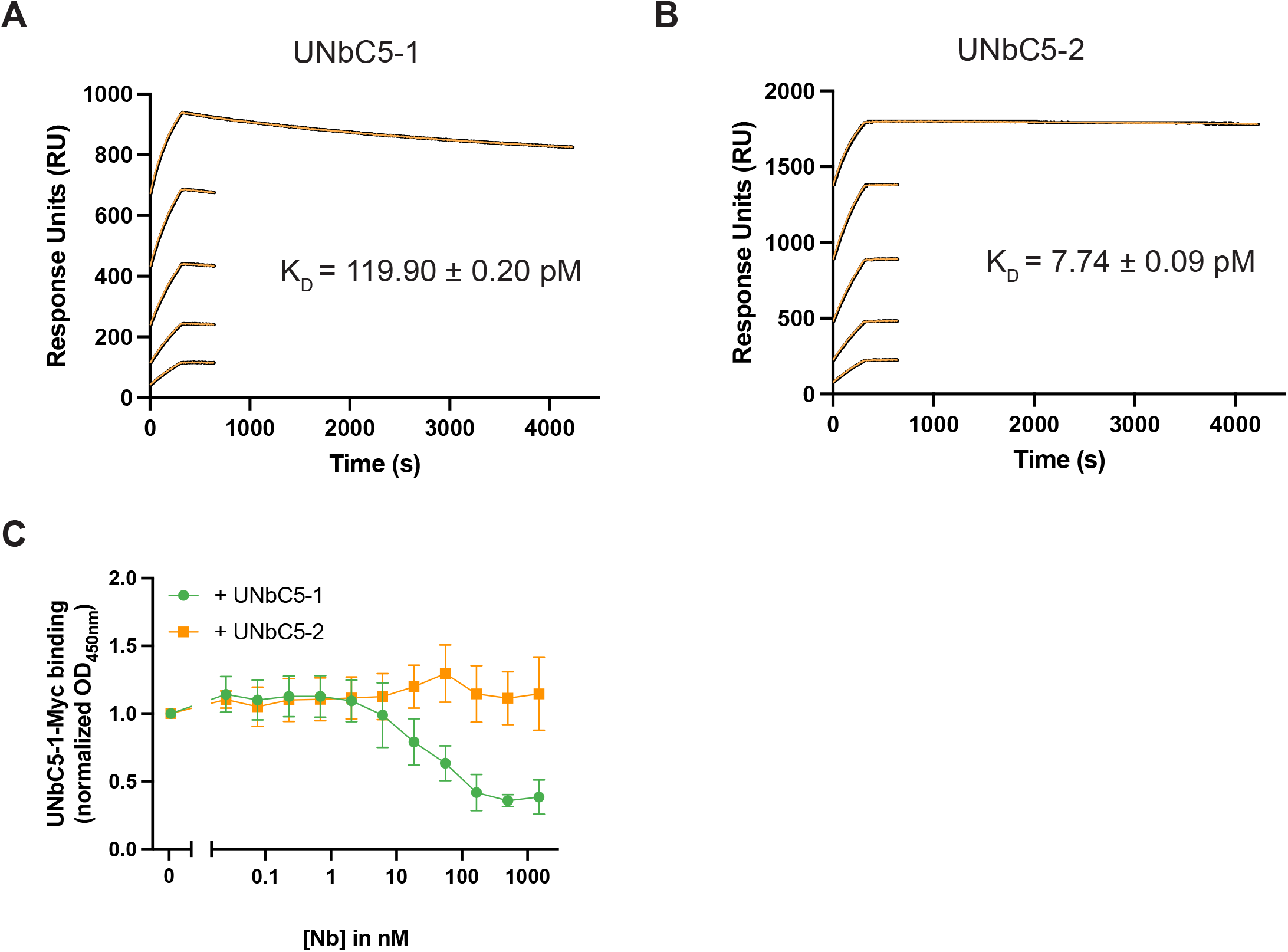
UNbC5-1 and UNbC5-2 bind C5 with picomolar affinity and recognize distinct epitopes. (A-B) SPR curves with UNbC5-1-biotin (biotinylation method [2]) (A) and UNbC5-2-biotin (biotinylation method [1]) (B) as a ligand and C5 as an analyte at concentrations of 12.5, 6.25, 3.13, 1.56, and 0.78 nM, evaluated over 4000 seconds. Experimental data is shown in black, and model fit in orange. (C) Competition ELISA with C5-coated microtiter plates incubated with 5 nM NbC5-1-Myc and a titration of untagged UNbC5-1 or UNbC5-2. Binding of UNbC5-1-Myc to C5 was measured using anti-Myc antibodies and an HRP-coupled secondary antibody, at OD450. Data was normalized on maximum binding of UNbC5-1-Myc, measured when no untagged nanobody was added. Data information: (A-B) Experiments were carried out in duplicate as individual experiments at 25°C. (C) Data represent mean ± SD of 3 individual experiments.

### Cryo-EM structures of UNbC5-1 and UNbC5-2 in complex with C5

To obtain structural insights into the nanobody binding sites and inhibitory mechanisms, we determined the structure of both nonoverlapping nanobodies in complex with C5, using single-particle cryo-EM. The micrographs collected showed well-distributed particles in vitreous ice (**SFig. 2A**). Image processing was performed in CryoSPARC v3.3/3, where the 2D classification already showed secondary structural features of C5 in complex with both nanobodies (**SFig. 2B**). 3D classification and refinement of the electron density map led to an overall map resolution of 3.6 Å, which allowed the subsequent model building of the C5:UNbC5-1:UNbC5-2 structure (**SFig. 2C**). The structure was built using the previously described C5 crystal structure (3CU7) and AlphaFold generated nanobody models (23, 24). The final refinement of the structure in Phenix 1.20.1 showed acceptable model statistics and stereochemistry (**Table III**). Due to a lack of density, the C-terminal C345c domain could not be modeled. A similar flexibility of this C-terminal domain was observed in other C5 cryo-EM structures (10, 23), and the different arrangement of C345c in different crystal structures (8, 11, 16, 23, 25).

**Table III.**
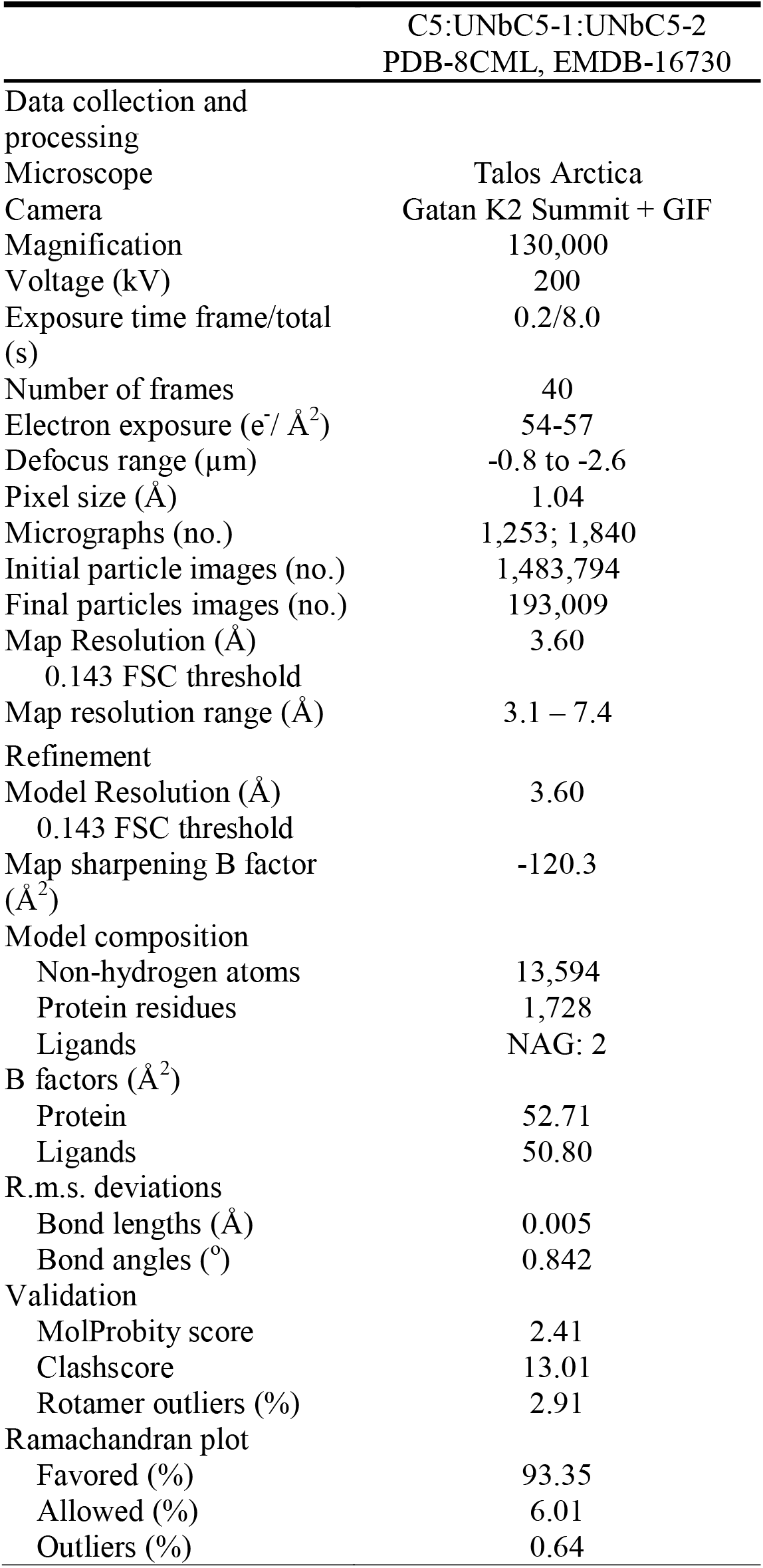
Cryo-EM data collection, refinement and validation statistics.

Overall, analysis of the C5 molecule in the C5:UNbC5-1:UNbC5-2 structure revealed interesting differences compared to the C5 crystal structure 3CU7 (**Fig. 3A**). Most apparent was the different arrangement of the C5a C-terminus, containing the C5 cleavage site, R751-L752. While residues D746-M754 are disordered in native C5 (3CU7), they adopt a helical conformation in the C5:UNbC5-1:UNbC5-2 structure, internalizing the cleavage site. This was also previously observed in C5 structures in complex with inhibitors Eculizumab-Fab, OmCI-RaCI3, knob K92, and NMR structures of C5a (11, 16, 25, 26). Superposition of MG1-MG6 domains from C5:UNbC5-1:UNbC5-2 and previously mentioned C5-inhibitor structures to C5 (3CU7) (**Fig. 3B, left panel**), show a similar C5a C-terminus rearrangement, likely preventing its cleavage by the C5 convertase (**Fig. 3B, right panel**) (11, 25). Additionally, all these structures show a similar rotational shift of the C5 α-chain, specifically domains C5a, CUB, MG8 and C5d, when compared to native C5 (3CU7). In conclusion, the differences between the structure of C5:UNbC5-1:UNbC5-2 and native C5 are similar to the previously described C5 structures in complex with known inhibitors.

**FIGURE 3:**
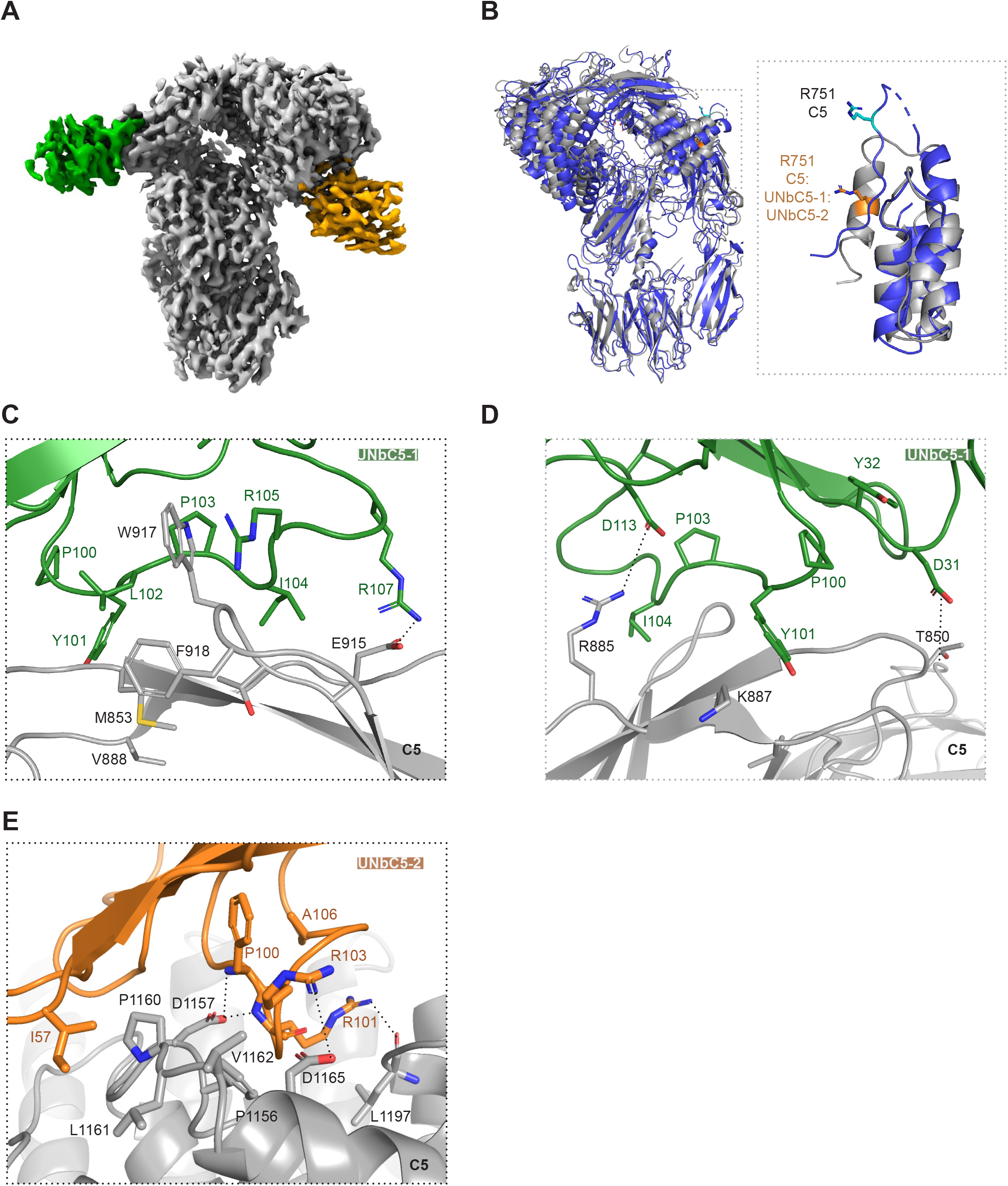
Cryo-EM structures of UNbC5-1 and UNbC5-2 in complex with C5. (A) Electron density map of the C5:UNbC5-1:UNbC5-2 complex at 3.6 Å resolution, with C5 colored in gray, UNbC5-1 in green, and UNbC5-2 in orange. (B) Comparison of the C5 structure excluding the C345c domain (3CU7, blue) to the C5 of the C5:UNbC5-1:UNbC5-2 structure (grey). Left panel shows both structures in cartoon representation, superimposed and aligned by the C5 MG-ring. Domains C5a, C5d, and CUB of the C5:UNbC5-1:UNbC5-2 structure shows a displacement when compared to the C5 structure. Right panel shows a zoom in of the domain C5a reoriented and aligned, both structures with residue R751 showed as sticks in orange for the C5:UNbC5-1:UNbC5-2 structure and in cyan for C5 (3CU7). (C-E) Zoom in on the C5 interface with nanobodies UNbC5-1 (C, D), and UNbC5-2 (E), in the same colors introduced in panel A. Amino acids that likely contribute to the interface are shown as sticks representation and annotated. Data information: (A) Electron density map visualized in ChimeraX and (B-F) figures produced in PyMOL from refined structure.

### Binding Interfaces

UNbC5-1 and UNbC5-2 bind to the C5 α-chain on opposite domains and with a different epitope-paratope architecture (**Fig. 3A**). While UNbC5-1 shows a more commonly observed nanobody/antibody interface consisting of a hydrophobic core surrounded by polar interactions, UNbC5-2 interface is mostly formed by salt bridges and hydrogen bonds.

C5 and UNbC5-1 buried a total interface of 1,535 Å^2^(27). UNbC5-1 binds to the MG7 domain, which is considered a known target for complement inhibition. For example, Eculizumab (and its derivatives) (16) and nanobodies targeting C5 homologues C4b (NbE3 and hC4Nb8) (18, 28) and C3/C3b (hC3Nb1) (29), bind to the MG7 domain. Proposed convertase models (SCIN-convertase (2WIN), and CVF-C5 (3PVM)) suggest the involvement of this domain in the convertase-substrate interaction (30, 31). UNbC5-1 binds to the connecting loops of the four-stranded antiparallel β-sheet in the MG7 domain (**Fig. 3C & D**). Most of the interface is formed by the extended CDR3, with limited contribution of CDR1 and 2. Residues Y101 to I104, of the CD3 loop, form a hydrophobic core that interacts with three of the MG7 loops with residues M853, V888, W917, and F918. Furthermore, R105 forms a cation-π interaction with residue W917. The paratope architecture is also maintained by residues Y32 of CDR1 and Y101 of CDR3 that sandwich P100 of CDR3, where Y101 interacts with MG7 residue K887. Additionally, CDR3 residues R107 and D113 form salt bridges with E915 and R885 of C5-MG7, respectively. Finally, CDR1 helps to stabilize the interface with interactions between CDR1 D31 and MG7 T850 and some hydrogen bonding contacts from CDR2 also contribute with the interface.

The C5:UNbC5-2 buried a surface area of 1,084 Å^2^ (27). The C5 binding epitope of UNbC5-2 comprises residues of several α-helices and loops of the C5d domain (**Fig. 3E**). Similar as the MG7 domain, the C5d domain is targeted by multiple complement inhibitors, including OmCI and RaCI3 (11). The C5 interface is formed by the three CDRs of the nanobody. Residue I57 of CDR2 interacts with a hydrophobic patch formed by residues P1160 to V1162. Additionally, some electrostatic interactions are formed between C5d residue K1091 and residue E99 of CDR3, and D1165 with R103 from CDR3. Moreover, D1157 of C5d forms hydrogen bonds with the backbone of residues P100 and R101 of CDR3, and D1157 backbone also interacts with residue T33 of CDR1. Other observed interactions are C5d residue Q1097 and residue N54 of CDR2 and staking of residue R103 with F1156 of C5d.

### The C5-binding interface of UNbC5-2 partly overlaps with RaCI3

The UNbC5-2 interface in C5 is close to that of RaCI3, therefore we next compared the structure of C5:UNbC5-1:UNbC5-2 with C5-OmCI-RaCI3 (5HCC) (**Fig. 4A**). RaCI3 binds to the cleft formed by domain MG1, MG2 and C5d, interacting with all 3 domains of C5 (11). A detailed comparison revealed six overlapping residues between the UNbC5-2 and RaCI3 interfaces on C5d (**Fig. 4B**) (11). In contrast to the C5-RaCI3 interactions, which are mostly driven by extensive van der Waals interactions and hydrogen bonds, the C5:UNbC5-2 interface include hydrophobic interactions with residues P1160-V1162, a hydrogen bond with Q1097 and a salt bridge with residue D1165 of C5d. To investigate whether UNbC5-2 and RaCI3 can bind C5 simultaneously, we performed an SPR assay. Here we used Eculizumab-Fab as a bait to capture C5 and sequentially injected RaCI3 and UNbC5-2. As expected, we measured association of C5 and RaCI3 (**Fig. 4C**). Interestingly, there was no increase in signal when UNbC5-2 was added. This indicates that UNbC5-2 and RaCI3 cannot bind C5 together. To conclude, the structural model and SPR assay indicate that the binding epitopes of UNbC5-2 and RaCI3 on C5 partially overlap and that both inhibitors cannot bind C5 simultaneously.

**FIGURE 4:**
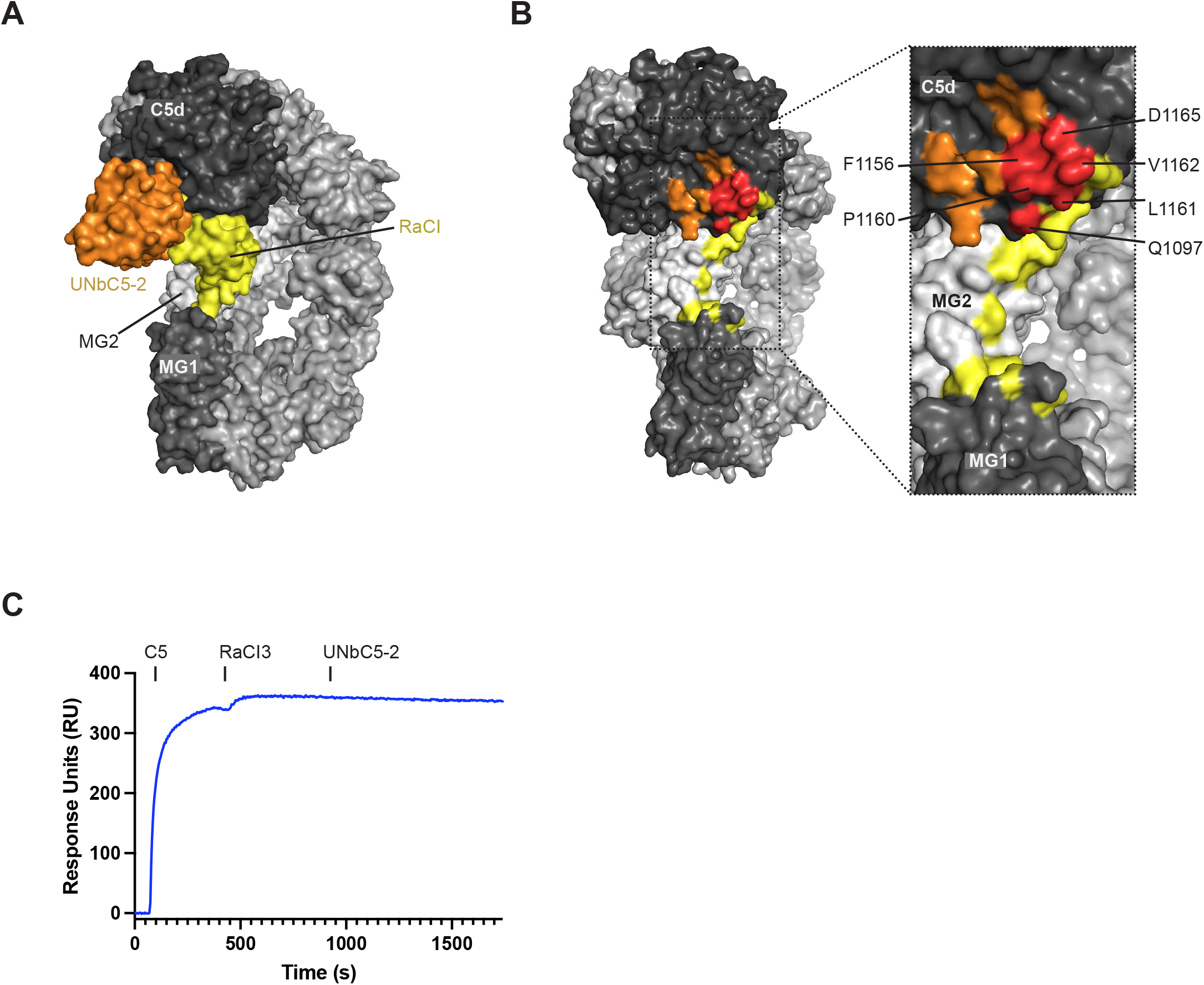
The C5-binding interface of UNbC5-2 partially overlaps with RaCI3. (A) Superposition of the C5:UNbC5-1:UNbC5-2 and the C5-RaCI3 complex in surface representation with UNbC5-2 in orange, RaCI3 in yellow, most of the C5 domains in light gray, MG1 in medium gray and C5d in dark gray. (B) Left panel shows a 45° rotation view of panel A, where footprint of epitopes of UNbC5-2, RaCI3 and their overlapping residues (red) are colored according to panel A. Right panel shows a zoom in of UNbC5-2 and RaCI3 epitopes where overlapping residues are annotated. (C) Competition ELISA with C5-coated microtiter plates incubated with 300 nM UNbC5-2 and a titration of RaCI3 or Eculizumab (5.49-4000 nM, 3-fold). Binding of UNbC5-2 to C5 was measured using polyclonal rabbit-anti-VHH QE19 antibodies and donkey-anti-rabbit-HRP antibodies, at OD450. (D) Competition SPR between UNbC5-2 and RaCI3. Eculizumab-Fab was used as a ligand, followed up by injections of 100nM C5, RaCI3 and UNbC5-2, at time points 60, 360, and 900 seconds, respectively. Injection of RaCI3 shows binding to C5 bound to Eculizumab-Fab. Additionally, sequential injection of UNbC5-2 at 900 seconds showed no signal increase, denoting competition. Data information: (A, B) Figures produced in PyMOL, (C) Data represent mean ± SD of 3 individual experiments.

### UNbC5-1 competes with Eculizumab on binding the MG7 domain of C5

A comparative analysis of C5-Eculizumab-Fab and C5:UNbC5-1:UNbC5-2, showed an extensive overlap of both the Eculizumab-Fab and UNbC5-1 epitope (**Fig. 5A**). The Eculizumab-Fab epitope extends through the posterior antiparallel β-strands of the MG7 domain, while the UNbC5-1 epitope is shifted downwards and binds both the β-strands and its connecting loops (16). Both paratopes interact through hydrophobic interactions with C5 residue W917 and form salt bridges with residues K887 and E915, in addition of other varied interactions with residues G852 and Q854 (**Fig. 5B**). Interestingly, they interact with R885 differently, UNbC5-1 forms a salt bridge with this residue, at the edge of the epitope interface, while in the Eculizumab-Fab interface the residue enters a hydrophobic pocket in the center of the interface formed by residues W33, F101 and W107. To confirm with an *in vitro* experiment that UNbC5-1 competes with Eculizumab for C5 binding, we performed a competition ELISA. Briefly, we coated C5 and measured binding of UNbC5-1 in the presence of a titration of Eculizumab. As a negative control we added RaCI3 instead of Eculizumab. Consistent with the structural data, we observed a decrease in UNbC5-1 binding in the presence of Eculizumab, but not with RaCI3 (**Fig. 5C**).

**FIGURE 5:**
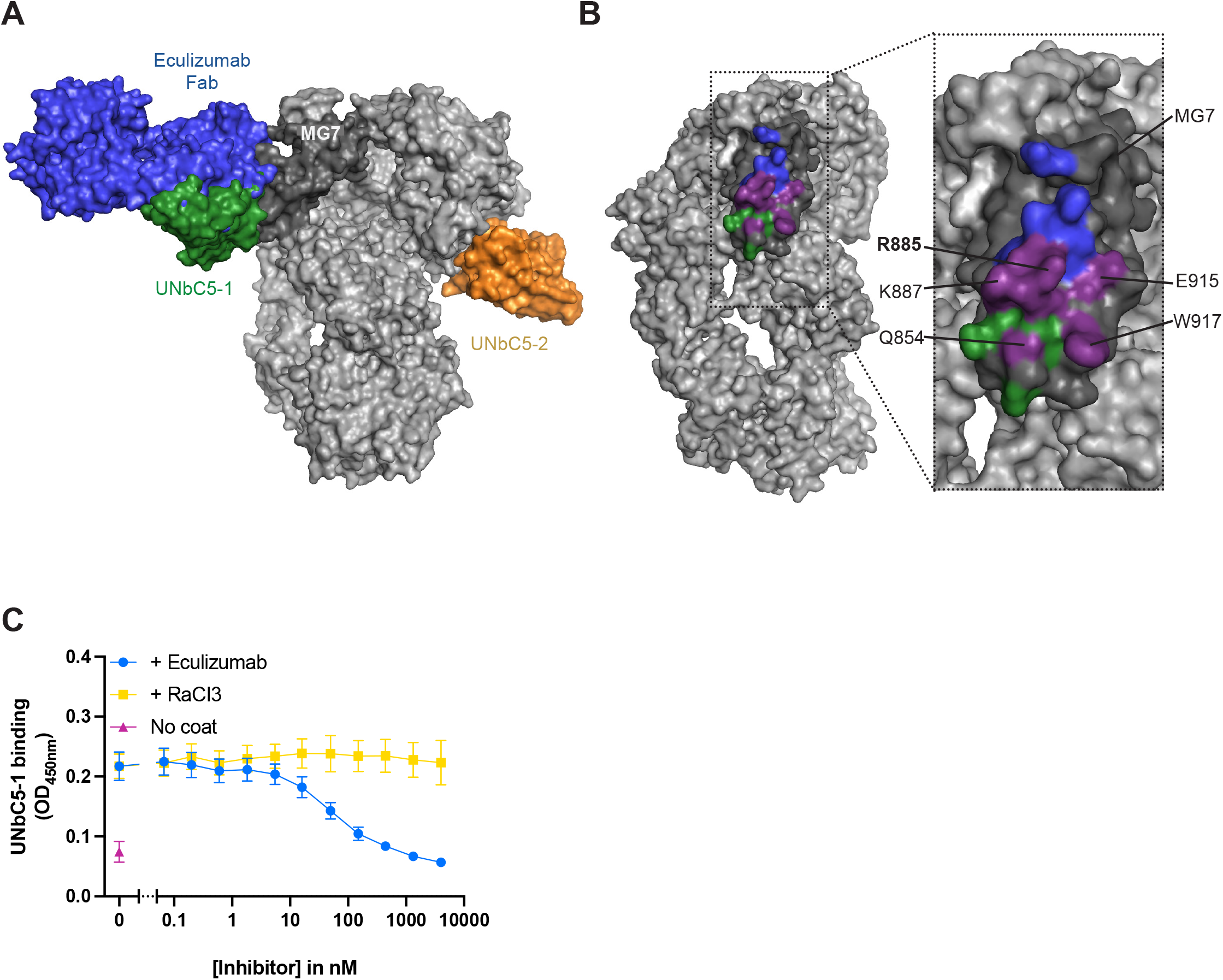
UNbC5-1 competes with Eculizumab in binding to the MG7 domain of C5. (A) Superposition of the C5:UNbC5-1:UNbC5-2 and the C5-Eculizumab-Fab complex structure. Surface representation shows epitope overlap between UNbC5-1 and Eculizumab-Fab. UNbC5-1 is shown in green, UNbC5-2 in orange, Eculizumab-Fab in blue, most domains of C5 in light gray and domain MG7 in dark gray. (B) Left panel shows a 45° rotation of panel A. Right panel shows a zoom in of the epitope of UNbC5-1 (green) and Eculizumab-Fab (blue) on the MG7 domain (dark grey) with an extensive overlap (purple) of the epitope between the interface of both molecules in C5. The overlapping amino acids for both interfaces are indicated, with residue R885 in bold. (C) Competition ELISA with C5-coated microtiter plates incubated with 300 nM UNbC5-1 and a titration of Eculizumab or RaCI3 (5.49-4000 nM, 3-fold). Binding of UNbC5-1 to C5 was measured using polyclonal rabbit-anti-VHH QE19 antibodies and donkey-anti-rabbit-HRP antibodies, at OD 450 nm. Data information: (A, B) Figures produced in PyMOL, (C) Data represent mean ± SD of 4 individual experiments.

### UNbC5-1 binds and inhibits the C5 variant R885H

Finally, we wondered if UNbC5-1 could recognize the genetic variant C5 R885H (15), that is not targeted by Eculizumab. First, we coated microtiter plates with C5 WT and C5 R885H, added UNbC5-1 or Eculizumab and measured their binding. As described before (15, 17), we observed in ELISA and SPR that Eculizumab binds C5 R885H poorly (**Fig. 6A & SFig. 4A**). Interestingly, UNbC5-1 binds C5 R885H efficiently (**Fig. 6B & Table I**), with a binding affinity of 15.5 nM (**SFig. 4B)**. This affinity is roughly 100× lower compared to the affinity for C5 WT, which confirms that residue 885 is involved in the binding interface of UNbC5-1 and C5, but is not crucial for its interaction, like it is for Eculizumab. Next, we assessed whether UNbC5-1 was also able to prevent complement-mediated erythrocyte lysis with C5 R885H. To assess this, we incubated antibody-coated sheep erythrocytes with C5 depleted serum, repleted with physiological concentrations of C5 WT or C5 R885H. Consistent with the binding data, we observed that Eculizumab failed to inhibit complement activity via C5 R885H (**Fig. 6C**). On the contrary, UNbC5-1 inhibits complement activity with C5 R885H and C5 WT to a similar extent (**Fig. 6D**). Finally, for UNbC5-2, which binds C5 to an epitope unrelated to amino acid 885, we also observed similar binding efficiencies in ELISA (**SFig. 4C)**. Finally, we show that UNbC5-2 inhibits complement mediated lysis with comparable potencies with C5 R885H and C5 WT (**SFig. 4D**). Altogether, these data show that UNbC5-1 and UNbC5-2 bind and inhibit different genetic variants of C5.

**FIGURE 6:**
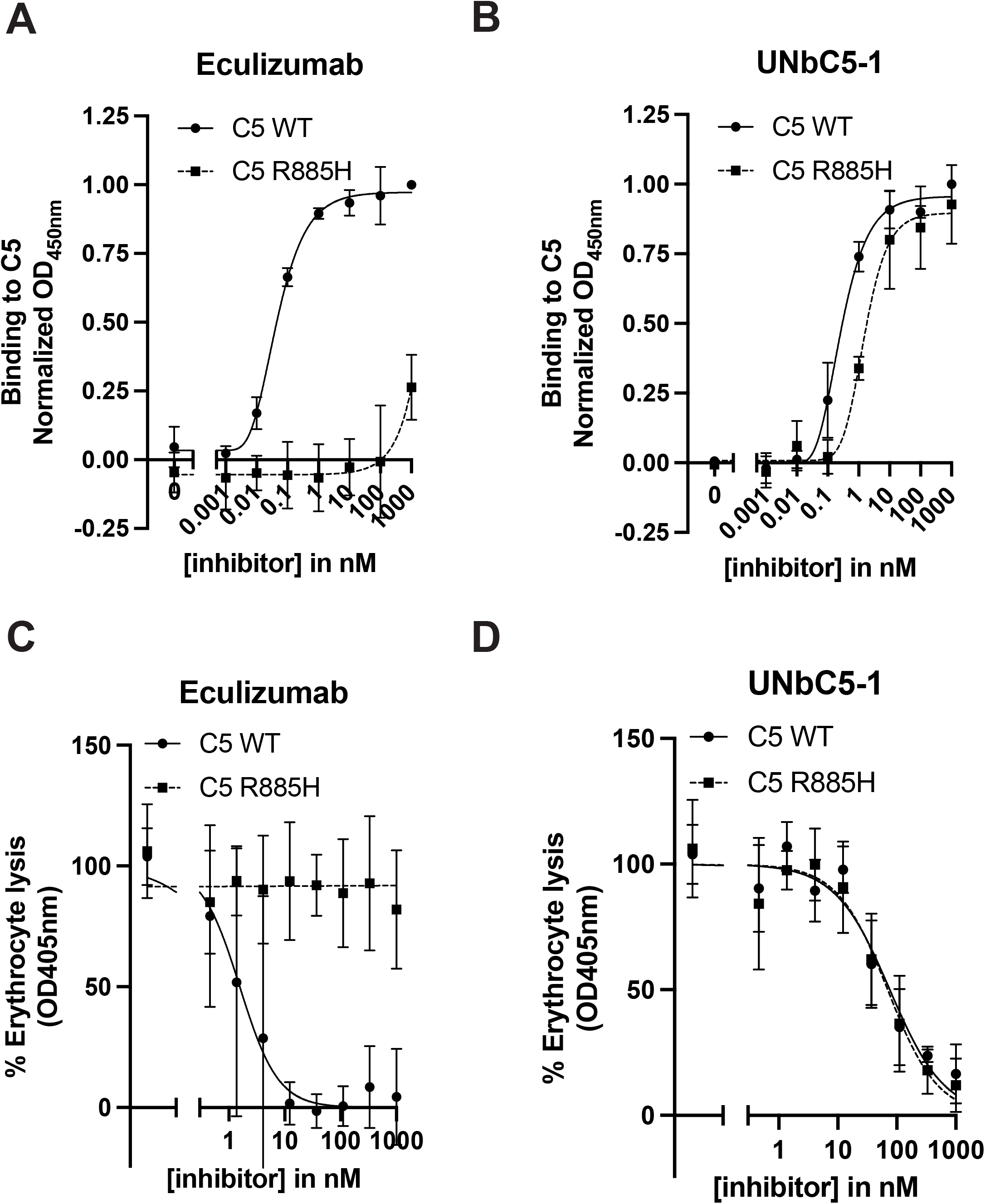
UNbC5-1 binds and inhibits C5 variant R885H. (A-B) Binding of Eculizumab (A) and UNbC5-1 (B) to C5 WT and C5 R885H, using C5 WT- or C5 R885H-coated microtiter plates, incubated with increasing concentrations of Eculizumab or UNbC5-1. Binding was assessed with a monoclonal anti-human-kappa antibody (A) or polyclonal rabbit-anti-VHH QE19 antibodies (B) and donkey-anti-rabbit-HRP secondary antibodies (A, B), at OD450. (C-D) CP mediated hemolysis of antibody-coated sheep erythrocytes incubated with 2.5% C5 depleted human serum, repleted with physiological concentrations of C5 WT or C5 R885H and a titration of Eculizumab (C) or UNbC5-1 (D). The OD450 values of the supernatants were measured, and the % erythrocyte lysis was calculated using a 0% (buffer) and 100% (MilliQ) control sample. Data information: (A-D) Data represent mean ± SD of 3 individual experiments and curves were fitted.

## Discussion

With the identification of UNbC5-1 and UNbC5-2 as specific, high affinity C5 targeting nanobodies, this study adds two new C5 inhibitors to the field. In the last decades, diverse C5 targeting molecules have been identified and developed, including monoclonal antibodies, single domain antibodies, small molecules, nucleic acid-based therapies, pathogen derived proteins and peptides, but no nanobodies (13). In contrast, nanobodies against other complement components (C1q (32), C3 (29, 33, 34), C4 (18, 35), properdin (36, 37)) and complement receptors (CRIg (38), Vsig4 (39)) were previously identified. The use of nanobodies as therapeutic molecules has gained increasing interest in many different fields, especially since the successful introduction of the first nanobody in the clinic, Caplacizumab (targeting plasma protein von Willebrand factor) (40). Compared to conventional antibodies, nanobodies lack an Fc-tail to activate the immune system, which is favorable when complement inhibition is the goal. At the same time, the lack of an Fc-tail shortens the half-life of nanobodies in circulation, while improving tissue penetration. In addition to their therapeutic potential, nanobodies are increasingly used as tools in research and diagnostics because they are easy to produce and label with for example fluorophores (41, 42) or to conjugate to solid matrixes for chromatography (43). Also, the presented C5-specific nanobodies could be developed for specific detection of C5 in complex biological specimens or for affinity purification of C5 from human plasma. Furthermore, their inhibitory action makes them suited as tools to inhibit the complement system at level of C5 cleavage, thereby studying the role of complement in diseases.

Elucidating the exact inhibitory mechanism by which our nanobodies prevent C5 cleavage is complicated by the lack of insight into how C5 interacts with C5 convertase enzymes. These multi-component complexes are unstable and exclusively formed on surfaces of biological organisms such as bacteria (44). This has hampered structural elucidation of the C5 convertase and its interaction with substrate C5. For UNbC5-1, it is likely that the inhibitory mechanism is similar to the clinically used Eculizumab because both structural and biochemical competition analyses showed that the nanobody binds to a similar interface as Eculizumab. For Eculizumab it was suggested that binding to the MG7 domain sterically hinders the interaction between C5 and the C5 convertase (16).

For UNbC5-2, the inhibitory mechanism is less evident. Because the binding interface of UNbC5-2 partially overlaps with RaCI3 (11), the inhibitory mechanism could be similar to RaCI. For the RaCI family it is suggested that binding to C5d (together with OmCI), induces a conformational change in which the normally disordered scissile loop (residues 746-754), near the cleavage site of C5a (R751-L752), is altered into an ordered α-helix. These conformational rearrangements lower the affinity of C5 for C3b and make the scissile loop unavailable for the C5 convertase (11, 16, 25, 45). Interestingly, the here presented structure of C5 with UNbC5-1 and UNbC5-2 strengthens the idea that C5 inhibitors from different sources can adopt a common inhibitory mechanism. As observed for Eculizumab-Fab (derived from mice (16)), OmCI-RaCI3 (derived from ticks (11)) and knob K92 (derived from cows (25)), our llama-derived nanobodies similarly induce a conformational change that leads to the formation of an ordered α-helical C5a C-terminus. Macpherson *et al*. already recognized that these domain movements could play a role in the Knob K92 inhibition mechanism (25). Our data suggest that this partial molecular movement of the α-chain and helical rearrangement of the scissile loop can be triggered by various inhibitors targeting different epitopes across the molecule (10, 16, 25). Because the α-helical arrangement is found in both the C5-OmCI-RaCI3 and C5-Eculizumab-Fab structure (11, 16), it is impossible with our data to attribute this proposed mechanism of inhibition to either UNbC5-1 or UNbC5-2, or both.

Although Eculizumab is proven to be an effective therapy in controlling diseases like aHUS, a drawback is therapy resistance in a small proportion of patients with C5 polymorphism (15). This was explained by the fact that the interaction of Eculizumab with the R885 residue is essential for binding. Interestingly, we observed that UNbC5-1 can bind and can inhibit the R885H variant of C5. Even though UNbC5-1 does interact with R885, the structure suggests that mutation of R885 into a histidine has less impact on the C5:UNbC5-1 interface, because R885 interacts mostly with a polar patch (formed by residues E915 of MG7, and R107 and D113 of CDR3 at the edge of the interface) and a H885 variant could probably still establish a polar interaction. In contrast, R885 extends into a hydrophobic pocket in the Eculizumab interface surrounded by bulky sidechain amino acids such as W33, F101, and W107 where a histidine sidechain might be too small to extend inward and interact, therefore disrupting the core interaction of the interface (16). We propose that the detailed binding interface of UNbC5-1 and C5 is a good starting point to further optimize reactivity of Eculizumab (and its Fc-optimized successor Ravulizumab) towards the genetic variant of C5 R885H.

To conclude, in this study we developed and characterized two anti-C5 nanobodies that bind C5 with picomolar affinities and efficiently inhibit complement at the level of C5 cleavage. Both nanobodies can be used to detect C5 and inhibit C5 cleavage for diagnostic and/or research purposes. Finally, both nanobodies have the potential to be further developed for therapeutic purposes and detailed binding interfaces obtained here could help to further develop high affinity C5 inhibitors that are reactive to different genetic variants of C5.

## Materials and methods

### Serum, proteins and complement inhibitors

Normal human serum was obtained from a pool of healthy donors as previously described (46). C5 depleted serum was obtained from Complement Technology. Complement protein C3 was isolated and purified from freshly obtained human plasma, as described before (47). Complement protein C4 was obtained from Complement Technology. Complement proteins C5 WT (unless stated differently) and C5 R885H and complement inhibitor RaCI3 were recombinantly produced and purified in our lab, using EXPI293F cells, similar to described for complement protein C9 (48), with a few adaptations. For C5 WT and C5 R885H the adaptations were: [1]C5 R885H DNA was created via overlap extension PCR, using the pcDNA-C5 vector as a template. [2] Supernatants after 5 days of expression were concentrated and buffer exchanged to 25 mM HEPES, 500 mM NaCl, 5 mM benzamidine, pH 8.2 using the QuixStand benchtop system (GE Healthcare) and 25 mM imidazole was added before application to the HiTrap Chelating column. [3] The HiTrap Chelating column was washed with 50 mM imidazole and protein was eluted with 175 mM imidazole. For RaCI3 we made the following adaptations: [1] C-terminal tags did not contain a 3×(GGGGS) linker. [2] 1.0 μg DNA/ml cells was used. [3] A HisTrap FF column (Cytiva, GE Healthcare) was used in the Äkta Pure protein chromatography system (GE Healthcare), according to manufacturer’s description. Before application to the column, the supernatant was dialyzed against 50 mM Tris, 500 mM NaCl, pH 8.0, and 30 mM imidazole was added. For C5 WT, C5 R885H and RaCI3 no additional purification step by size exclusion chromatography was performed and proteins were dialyzed to PBS and stored at -80°C directly after the HiTrap Chelating column. Complement inhibitor Eculizumab was produced and purified in our lab. Briefly, the heavy and light chain variable plus constant region amino acid sequences of Eculizumab (14) were from patent (WO2017044811A1). gBlocks containing codon optimized sequences with an upstream KOZAK and HAVT20 signal peptide were cloned into the pcDNA34 vector, using Gibson assembly (New England Biolabs). After verification of the correct sequence, plasmids were used to transfect EXPI293F cells (Thermo Fisher Scientific) using 1 μg DNA/ml of cells with a ratio of light chain:heavy chain of 3:2. Expression and purification of Eculizumab was performed similar to described for complement protein C9 (48) with the added requirement that >95% of Eculizumab appeared as monomeric IgG.

### Nanobody production and purification

#### Nanobodies with Myc- and His-tags

Nanobody genes were obtained from the mRNA isolated from llama-derived PBMCs and after multiple PCR steps cloned into a pPQ81 vector, a derivative of pHEN1 (49). This provided C-terminal Myc- and His-tags, and nanobodies were expressed in *E. coli* Rosetta 2 (DE3) BL21 cells (Merck). Transformed bacteria were inoculated in 2× Yeast Extract Tryptone (2YT) medium supplemented with 2% glucose and 100 μg/ml ampicillin and grown shaking, at 37°C, overnight (O/N). The next day cultures were diluted 1:20 in 2YT medium supplemented with 0.1% glucose and 100 μg/ml ampicillin and grown at 37°C, shaking, until an OD_600_ of 0.6-0.9 was obtained. Then, nanobody production was induced with 1 mM isopropylthio-β-galactoside and expression was continued for 4 hours at 37°C or O/N at room temperature (RT), shaking. Next, cultures were spun down for 10 minutes at 6000 g. Pellets were dissolved in PBS and frozen for >30 minutes at -80°C or O/N at -20°C and thawed, to release periplasmic fractions in the supernatant. Thawed pellets were centrifuged for 15 minutes at 6000 g. Next, immobilized metal-affinity chromatography (IMAC) was used with ROTI Garose-His/Co beads (Roth) to purify nanobodies via their His-tag. After binding to the beads, nanobodies were eluted with 150 mM imidazole. Subsequently, nanobodies were dialyzed to PBS and concentrations were determined at OD_280_ using the Nanodrop One (Thermo Fisher Scientific). Finally, nanobody size, purity and concentration were verified using SDS-Page, stained with InstantBlue Safe Coomassie stain (Sigma Aldrich).

#### Nanobodies without a Myc-tag

To produce nanobodies without a Myc-tag, nanobody genes were recloned in the pYQ11 vector (previously published as pYQVQ11 (50)), which encodes for a C-terminal C-Direct tag containing a free thiol (cysteine) and an EPEA (Glu, Pro, Glu, Ala) purification tag (C-tag, Thermo Fisher Scientific). Vectors were transformed into yeast cells (Saccharomyces cerevisiae strain VWK18 (51)). For expression cells were inoculated in Yeast Peptone (YP) medium supplemented with 2% glucose and grown for 24 hours, at 30°C, shaking. Next, cultures were diluted 1:20 in Yeast Nitrogen Base (YNB) medium supplemented with 2% glucose and grown at 30°C, shaking. After 24 hours, cultures were diluted 1:10 by adding YP medium, supplemented with 2% glucose and 1% galactose. Nanobody production was continued for >64 hours, at 30°C, shaking, until an OD_600_ of >20 was reached. Next, cells were spun down for 20 minutes at 6000 g and supernatants were collected, filter sterilized (0.45 μm) and stored at 4°C until purification. Produced nanobodies were purified from yeast supernatants using a CaptureSelect C-tag column (Thermo Fisher Scientific) and an Äkta Start (Cytiva). For this, the supernatants were adjusted to neutral pH by adding 10× PBS. Bound nanobodies were washed with at least 5 column volumes of PBS and eluted off with 20 mM citrate buffer supplemented with 150 mM NaCl, pH 3.0. Next, samples were neutralized with 100 mM Tris, pH 7.5 and dialyzed to PBS. Protein concentration was determined at OD_280_ using a MultiScan (Thermo Fisher Scientific). Finally, nanobody size, purity and concentration were confirmed using SDS-PAGE, followed by Instant Blue Safe Coomassie staining.

#### Biotinylated nanobodies and Eculizumab(-Fab)

Nanobodies and Eculizumab(-Fab) were biotinylated using two methods. [1] Nanobody sequences were modified to contain a C-terminal LPETG-His motif and were subsequent produced and purified in *E. coli* BL21 Rosetta, like described above. After purification 50 μM LPETG-His nanobody was incubated with 25 μM His-tagged Sortase A7 (His-Tevg-SrtA7) (52) and 1 mM GGGK-Biotin (provided by Louris Feitsma, Medicinal Chemistry, Utrecht University), in 50 mM Tris, supplemented with 300 mM NaCl, for 2 hours, at 4°C. His-Tevg-SrtA7 was modified, expressed in *E. coli* and purified using its His-tag, in our lab. Next, samples were equilibrated with 20 mM imidazole and run over an HisTrap FF column to remove all His-tagged compounds from the reaction. Subsequently, a 30 kDa Amicon Tube (Merck Millipore) was used to concentrate the biotinylated nanobodies. To remove excess GGGK-biotin, samples were run over a Zeba™ Spin Desalting column (Thermo Fisher Scientific). [2] Myc-His-tagged nanobodies produced in *E. coli* and Eculizumab antibody and Eculizumab-Fab, were randomly biotinylated via N-hydrocysucciminimidyl (NHS)-labeling. Briefly, nanobodies at a concentration of 50-70 μM were incubated for 2 hours, at 4°C, with 20× molar excess of EZ link™ NHS-PEG4-Biotin (Thermo Fisher Scientific, 210901BID), in PBS, pH 7.4. Unreacted linker was separated with Bio-Spin 6 columns (BioRad), that were previously equilibrated in PBS. The nanobody concentrations were determined using a Nanodrop One.

### Erythrocyte lysis assays

Sheep (Alsever Biotrading) and rabbit blood (kindly provided by Utrecht University, Faculty of Veterinary Medicine) was washed 3× with PBS and diluted to a 2% erythrocyte suspension in VBS (Veronal buffered saline + 145 nM NaCl, pH7.4). For the CP, sheep erythrocytes, pre-opsonized with 1:2000 polyclonal rabbit-anti-sheep IgM (produced in our lab), were used and VBS was supplemented with 0.25 mM MgCl_2_ and 0.5 mM CaCl_2_ (VBS++). For the AP, rabbit erythrocytes were used and VBS was supplemented with 5 mM MgCl_2_ and 10 mM EGTA (VBS/MgEGTA). Nanobodies were titrated (0.45-1000 nM, 3-fold) and incubated for 10 minutes at RT with normal human serum (CP: 2.5%, AP: 10%) or 2.5% C5 depleted serum, repleted with physiological concentrations of C5 WT or C5 R885H. Next nanobody-serum mixes were added to 2% erythrocyte suspensions and samples were incubated at 37°C, shaking, (CP: 10 minutes, AP: 30 minutes). Afterwards plates were spun down for 7 minutes at 3500 rpm. Supernatants were diluted 1:3 with MilliQ and haemoglobin release was measured in a flat-bottom plate, using an iMark Microplate Reader (Biorad) at 405 nm.

### ELISA: Coating, incubation times & development

Unless stated differently, Nunc Maxisorp plates (VWR 735-0083, Thermo Fisher Scientific) were coated with 2 μg/ml purified protein, diluted in PBS, in a volume of 50 μl. Plates were incubated O/N, at 4°C, non-shaking. Next, plates were blocked with 80 μl 4% BSA in PBS + 0.05% Tween-20 (PBS-T). All following incubation steps were performed in 50 μl 1% BSA PBS-T, shaking, at RT, for 1 hour. In-between all steps, plates were washed 3× with PBS-T. To develop the ELISAs, tetramethylbenzidine (TMB) substrate solution of 6 mg/ml, dissolved in DMSO, was used to activate the enzyme labeled antibodies. When a color change was observed, the reaction was stopped with 0.5 M sulfuric acid and absorbance was measured at 450 nm using an iMark Microplate Reader.

### Binding ELISA

Wells coated with purified complement components C3, C4, C5 WT or C5 R885H were used to assess the binding of our nanobodies and Eculizumab to different complement proteins. Nanobodies and Eculizumab were added in a 10-fold titration (0.001-1000 nM). Next, wells containing nanobodies were incubated with 1:2000 primary polyclonal rabbit-anti-VHH antibody QE19 (QVQ Holding BV) and 1:5000 secondary polyclonal donkey-anti-rabbit-HRP antibodies (Jackson Immuno Research). To detect Eculizumab binding, wells were incubated with anti-human-kappa-HRP (Southern Biotech) detection antibodies.

### Competition ELISA

Plates coated with purified complement component C5, were used to assess if our nanobodies compete. 5 nM UNbC5-1-Myc was incubated with a titration of untagged UNbC5-2 (0.025-1500 nM, 3-fold) for 10 minutes, at RT and subsequently added to the C5 coated wells. Next, wells were incubated with 1:2000 primary monoclonal mouse-anti-Myc-tag clone 9B11 #2274 (Cell Signaling Technologies), followed by 1:5000 secondary monoclonal goat-anti-mouse-PO (Southern Biotech). To assess competition between our nanobodies and Eculizumab/RaCI3, 300 nM of UNbC5-1 or UNbC5-2 was incubated with a titration of Eculizumab or RaCI3 (5.49-4000 nM, 3-fold) for 10 minutes at RT, prior to adding it to the C5 coated wells. Next, nanobody binding was detected using primary polyclonal rabbit-anti-VHH antibody QE19 and 1:5000 secondary polyclonal donkey-anti-rabbit-HRP antibodies.

### Classical Pathway Complement ELISA

Plates were coated with 3 μg/ml human IgM (Millipore) in 0.1 M sodiumcarbonate, pH 9.6. 1000 nM inhibitor (UNbC5-1, UNbC5-2, Eculizumab or RaCI3) was incubated with 4% normal human serum, added to the IgM coated wells, and incubated for 1 hour, at 37°C, shaking. To measure C5a formation, 25 μl of the supernatant was diluted with 25 μl 1% BSA PBS-T and added to a Nunc Maxisorp plate coated with 1 μg/ml anti-C5a capture antibody (C5a DuoSet ELISA kit, R&D systems). Next, we detected C5a with 2 μg/ml anti-C5a detection antibody (C5a DuoSet ELISA kit, R&D systems), followed by an incubation of 1:5000 streptavidin-HRP (Southern Biotech). To measure deposition of C3b on the IgM coated plates, 1:10,000 primary anti-C3 WM-1-DIG labeled antibodies (produced in our lab), and 1:8000 secondary anti-DIG-PO antibodies (Roche) were added. To measure deposition of C5b-9 on the IgM coated plates, 1:1000 primary monoclonal mouse-anti-C5b-9 aE11 (produced in our lab) and 1:5000 secondary polyclonal goat-anti-mouse-PO were used.

### Surface plasmon resonance (SPR)

Planar streptavidin coated chips (P-strep, Sens BV) were spotted with biotinylated UNbC5-1 (method [2]), UNbC5-2 (method [1]), and Eculizumab (method [1]) (25 and 50 nM, in duplicate) under a continuous flow for 1 hour using a microspotter (Wasatch). Subsequent SPR experiments were carried out in the IBIS-MX96 (IBIS Technologies) with SPR buffer (20 mM HEPES pH 7.4, 150 mM NaCl, 0.005% Tween-20). Initial testing showed limited regeneration with different high ionic strength (20 mM HEPES, 2 M NaCl, pH 7.4); ion containing (20 mM HEPES, 150 mM NaCl, 1 M MgCl_2_, pH 7.4); or low pH regeneration buffers (10 mM Glycine, pH 3.0). Therefore, the method of kinetic titration was selected to perform accurate affinity determination. C5 was injected in a series of 14 steps 2-fold dilutions, at concentrations of 0.78, 1.56, 3.13, 6.25, and 12.50 nM, without regeneration on nanobodies or Eculizumab coated surfaces. Determination of affinities for C5 R885H were performed with the same protocol, using concentrations of 25, 50, 100, 200, and 400 nM. The last step of dissociation ran for 60 and 35 minutes, respectively, for reliable determination of the K_off_. Kinetics were determined using Scrubber 2.0 (BioLogic Software), where simple bimolecular models were used to fit the data. Due to unsuccessful coating on the chip surface and the instability of RaCI3 and C5, we used Eculizumab-Fab-biotin (method [2]) as a ligand in the competition assay. C5, RaCI3 and UNbC5-2 were injected sequentially at concentrations of 100 nM, with 5 minutes association and 4 minutes dissociation for RaCI3 and UNbC5-2.

### Cryo-electron microscopy

#### Sample Preparation and Data Collection

C5, purchased from Complement Technologies, was diluted to a final concentration of 1 μM and gently mixed with a 1.5× molar excess of UNbC5-1 and UNbC5-2. The sample was diluted in PBS and incubated on ice for 20 minutes before freezing on glow discharged R1.2/1.3 200 mesh Au holey carbon grids (Quantifoil). The grids were then plunge frozen in ethane using a Vitrobot Mark IV (Thermo Fisher Scientific), at 4°C. Cryo-EM data was collected on a 200 kV Talos Arctica microscope (Thermo Fisher Scientific) equipped with a K2 summit detector (Gatan) and a post column 20 eV energy filter. Movies were collected in EPU (Thermo Fisher Scientific) at a magnification of 130,000× with a pixel size of 1.04 Å/pix. Two datasets were collected in two collection sessions with similar settings of 40 frames with a total exposure of 54 and 57 e-/Å^2^, and a defocus range of -0.8-2.6 μm.

#### Data Processing

The datasets of the C5:UNbC5-1:UNbC5-2 complex contained 1461 and 2349 movies (SFig. 2A). Both data sets were processed independently in CryoSPARC v3.2/3 with the same workflow until 2D classification. Overall, the workflow for each dataset started with patch-based motion correction and patch-based contrast function estimation (CTF) in CryoSPARC, followed by exposure curation that led to a total of 1253 and 1840 exposures, respectively. Then 100 movies of each data set were selected to perform a round of blob picking. Particles were extracted three times binned to 3.14 Å/pix and several rounds of 2D classification were performed to clean and select adequate templates for the CryoSPARC template picker. After template selection, the particles were picked and extracted unbinned with a 320-pixel box, particles of both datasets were merged, and an initial cleanup was performed through 2D classification. 1,483,794 particles from 2D selection were then submitted to generate four initial models (SFig.2B), that resulted in one good model that was further submitted to two initial model rounds (SFig. 2C). The resulted model with 193,009 particles was further selected for refinement. One round of non-uniform refinement led to an electron density map with a 3.73 Å resolution, which was further improved to 3.60 Å after local and global CTF refinement followed by a final non-uniform refinement. Global resolution was calculated according to the gold standard FSC = 0.143 criterion (SFig. 2D). Postprocessing such as sharpening and local resolution was also performed in CryoSPARC (SFig. 2C & E).

#### Model building and refinement

To build the model of the C5:UNbC5-1:UNbC5-2 complex, models for the two nanobodies were generated through AlphaFold (24) based on nanobody sequences. The C5 molecule comprising chains α and β of PDB 3CU7 (23), was also used in combination with the nanobody models to rigid-body fit them into the cryo-EM maps using UCSF ChimeraX (53, 54). The model was then iteratively refined using Coot and Phenix real-space refine (55, 56) with geometric restraints. The C345c domain of C5 was not included in the final structures because of the weak density in the map.

#### Data analysis & statistical testing

Nanobody sequences were aligned using T-coffee (57). Bar and line graphs were created using GraphPad Prism 9.3.0. Binding and inhibition curves were fitted in GraphPad Prism 9.3.0 using the functions “[inhibitor] vs. normalized response -- Variable slope” & “Asymmetric Sigmoidal, 5PL, X is concentration”. Fitted curves of 3 individual experiments were used to calculate IC50 and EC50 values with SD in GraphPad Prism 9.3.0. Figures with protein structures were prepared using PyMOL (retrieved from: http://www.pymol.org/pymol) (27). Figures with cryo-EM densities were created using UCSF ChimeraX (53, 54) The statistics of the structure come from the refinement in Phenix, SPR from Scrubber 2.0.

### Data availability

The model coordinates and cryo-EM density maps of the C5 structure in complex with nanobodies UNbC5-1 and UNbC5-2 have been deposited under the following accession numbers 8CML and EMD-16730.

## Supporting information

This article contains supporting information.

## Acknowledgments

We thank Wouter Beugelink and Dr. Itziar Serna Martin for their valuable input and scientific advice.

## Funding and additional information

This work was mainly supported by the Netherlands Organisation for Scientific Research (NWO) under the TTW Industrial Doctorate (grant agreement no. NWA.ID.17.036 to EMS) and the Consejo Nacional de Ciencia y Tecnología in Mexico (grant agreement No. CVU 604718 to KdB). The project also received funding from the European Research Council (ERC) under the European Union’s Horizon 2020 research and innovation programme (grant agreement No. 101001937, ERC-ACCENT to SHMR and No. 787241, ERC-C-CLEAR to PG).

## Conflict of interest statement

ED and RH are employees of QVQ Holding BV. Other authors declare no conflict of interest.

## Abbreviations

The abbreviations used here are:

sdAb: single-domain antibody
SPR: surface plasmon resonance
C5b-9 & MAC: membrane attack complex
PNH: paroxysmal nocturnal hemoglobinuria
aHUS: atypical hemolytic uremic syndrome
AMD: age-related macular degeneration
SLE: systemic lupus erythematosus
CP: classical pathway
LP: lectin pathway
AP: alternative pathway
CDR: complementarity determining regions
2YT: 2× yeast extract tryptone
O/N: overnight
RT: room temperature
IMAC: immobilized metal-affinity chromatography
YP: yeast peptone
YNB: yeast nitrogen base
NHS: N-hydrocysucciminidyl
VBS: veronal buffered saline
PBS-T: phosphate buffered saline supplemented with Tween
TMB: tetramethylbenzidine
ERC: European Research Council

