## Supplementary figures 1-4 for "Inhibition of cleavage of human complement component C5 and the R885H C5 variant by two distinct high affinity anti-C5 nanobodies"

**Material included:**
- 4 supporting figures

### Supporting Figure 1: Inhibition of the alternative pathway by UNbC5-1 and UNbC5-2

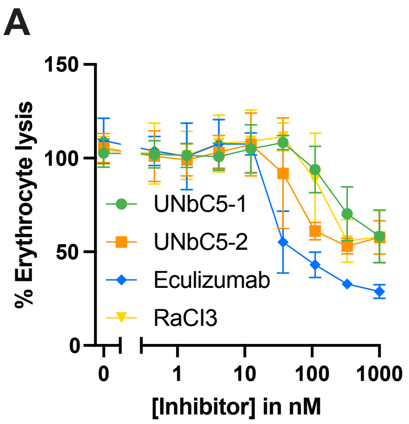


**SUPPLEMENTARY FIGURE 1: Inhibition of the alternative pathway by UNbC5-1 and UNbC5-2.** (A) AP mediated hemolysis of rabbit erythrocytes incubated with 10% normal human serum and a titration of our nanobodies UNbC5-1 and UNbC5-2 and known complement inhibitors RaCI3 and Eculizumab. The OD405 values of the supernatants were measured and the % erythrocyte lysis was calculated using a 0% (buffer) and 100% (MilliQ) control sample. Data information: (A) Data represent mean ± SD of 3 individual experiments.

**Supporting Figure 2: Cryo-EM image processing of C5 in complex with UNbC5-1 and UNbC5-2 datasets**

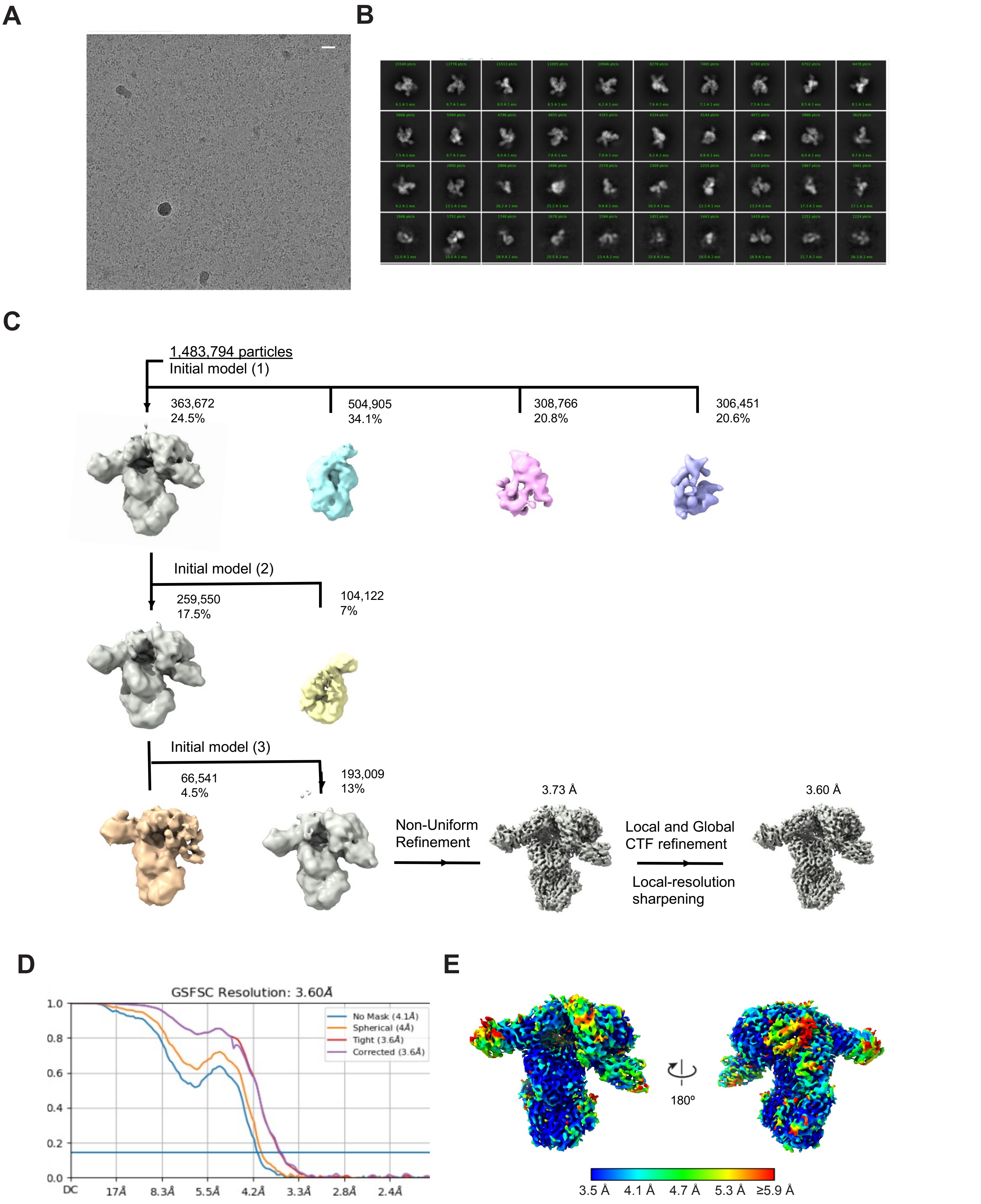


**SUPPLEMENTARY FIGURE 2: Cryo-EM image processing of C5 in complex with UNbC5-1 and UNbC5-2 datasets.** (A) Micrograph of C5:UNbC5-1:UNbC5-2 particles in vitreous ice. The scale bar length is 200 Å. (B) Selected 2D-class averages of the complex generated in CryoSPARC. (C) Three rounds of 3D Initial model classification and subsequent refinement strategy for the C5:UNbC5-1:UNbC5-2 reconstruction. (D) Fourier-shell correlation plots for the gold-standard refined C5:UNbC5-1:UNbC5-2 reconstruction, computed from unmasked (blue), with a spherical mask (orange), tight mask (red), and corrected (purple). (E) C5:UNbC5-1:UNbC5-2 complex density map colored by local resolution (blue = high, red = low), computed in CryoSPARC.

**Supporting Figure 3: Determination of the affinity of Eculizumab for C5**

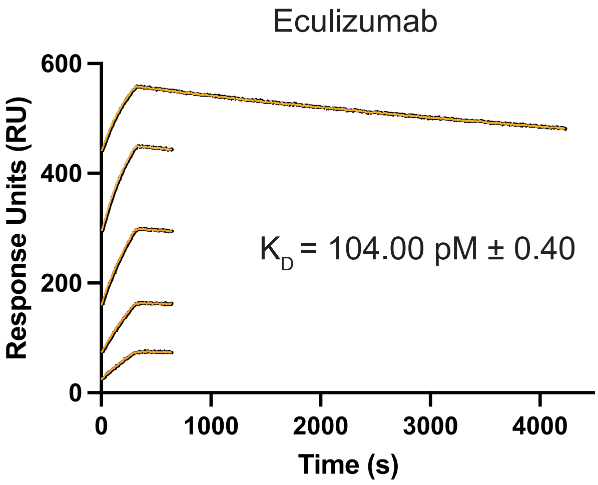


**SUPPLEMENTARY FIGURE 3: Determination of the affinity of Eculizumab for C5.** SPR curves of Eculizumab as a ligand and C5 as an analyte at concentrations of 12.5, 6.25, 3.13, 1.56, and 0.78 nM evaluated over 4000 seconds.

**Supporting Figure 4: Affinity determination for UNbC5-1 and Eculizumab for C5 R885H and UNbC5-2 binding to and ihibition of C5 R885H**

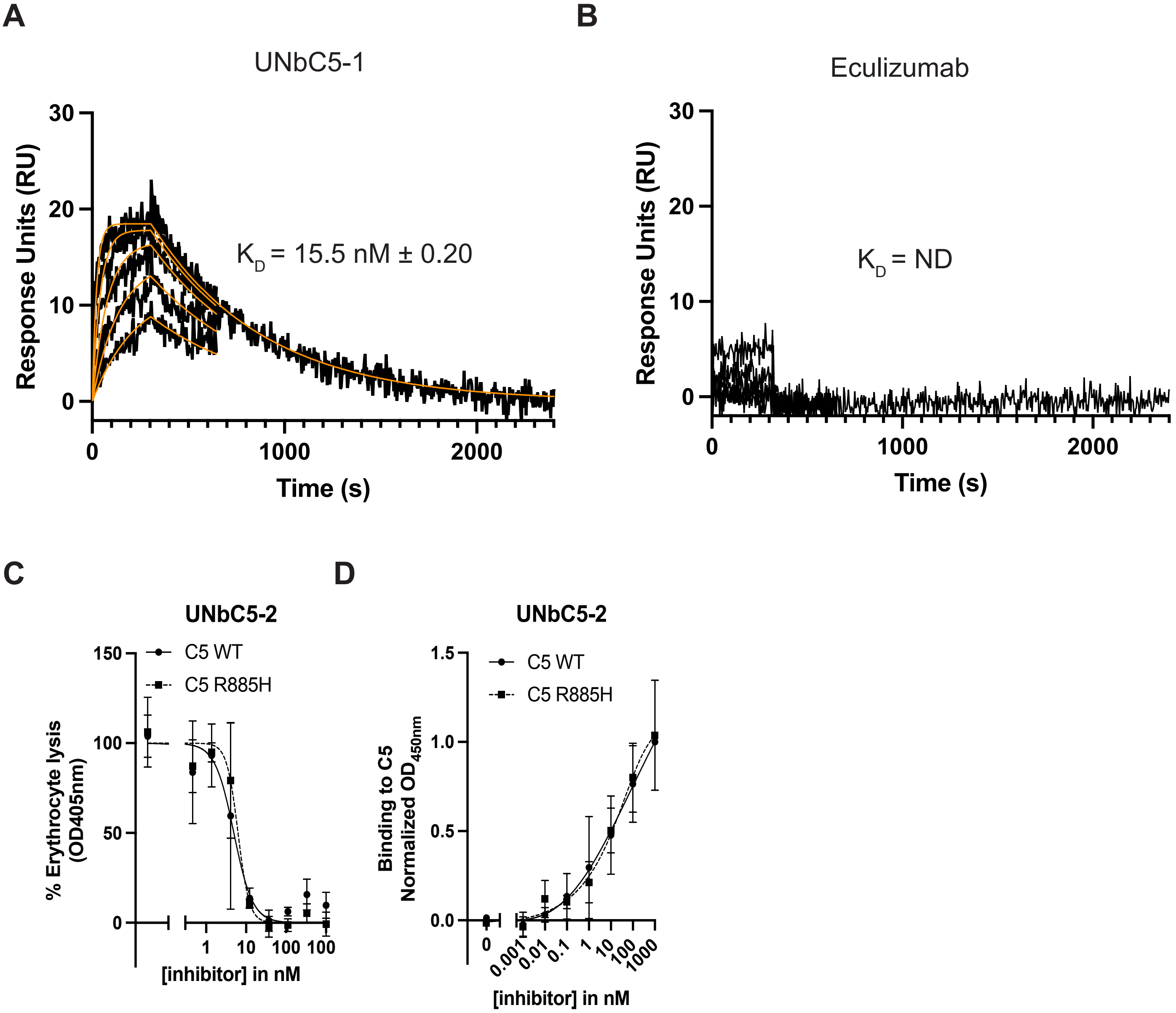


**SUPPLEMENTARY FIGURE 4: UNbC5-2 binding to and inhibition of C5 R885H.** (A-B) SPR curves with UNbC5-1 (A) and Eculizumab (B) as a ligand and C5 R885H as an analyte at concentrations of 400, 200, 100, 50, and 25 nM evaluated over 2400 seconds. (C) Binding of UNbC5-2 to C5 WT and C5 R885H, using C5 WT or C5 R885H coated microtiter plates, incubated with increasing concentrations of UNbC5-2. Binding was assessed with a monoclonal anti-human-kappa antibody and an HRP-labeled secondary antibodies, at OD450. (D) CP mediated hemolysis of antibody-coated sheep erythrocytes incubated with 2.5% C5 depleted human serum, repleted with physiological concentrations of C5 WT or C5 R885H and a titration of UNbC5-2. The OD405 values of the supernatants were measured, and the % erythrocyte lysis was calculated using a 0% (buffer) and 100% (MilliQ) control sample. Data represent mean ± SD of 2 (A-B) and 3 (C-D) individual experiments. (C-D) Curves were fitted.
